# RIFRMap: an interactive platform for integrating and visualizing rifampicin resistance mutations across bacteria

**DOI:** 10.64898/2026.11.28.769853

**Authors:** Lê Na Ngô, Andrew D. Letten, Jan Engelstädter

## Abstract

Rifampicin is among the most clinically important antibiotics in use today, yet resistance to it remains a global health burden. Although a growing body of work has illuminated the molecular basis of rifampicin resistance, predominantly mutations in the *rpoB* gene, the relevant data are dispersed across heterogeneous databases with different taxonomic scopes and curation standards, making cross-species comparison difficult. To address this, we developed RIFRMap (Rifampicin Resistance Mutation Map), an R Shiny web application that consolidates *rpoB* resistance mutation data from multiple sources. The current version comprises 1586 records covering 394 mutations across 81 bacterial species. Every position is mapped to the *Escherichia coli* MG1655 reference sequence so that organisms and datasets can be compared directly. The application provides interactive filtering and visualization modules that help users to explore mutation distributions, identify resistance hotspots, and assess consistency across datasets of rifampicin resistance. By unifying disparate resources into an interactive visual platform, RIFRMap provides researchers, clinicians, and educators with a means to conduct comparative analyses and investigate the global molecular landscape of rifampicin resistance.

## Introduction

Rifampicin (RIF), also known as rifampin, is one of the most clinically important broad-spectrum antibiotics and a first-line treatment for tuberculosis (TB). In addition to TB, RIF is used in the treatment of a range of other bacterial infections, including leprosy (1), pneumococcal meningitis (2), pulmonary disease caused by *Mycobacterium kansasii* (3), and infections caused by *Staphylococcus* and *Legionella* species (4, 5). The antibiotic exerts its bactericidal effect by binding to the *β* subunit of bacterial RNA polymerase (RNAP), encoded by the *rpoB* gene, thus blocking the extension of the nascent RNA chain (6). Given its clinical importance, resistance to RIF constitutes a major public health threat. According to the World Health Organization (WHO), approximately 370000 cases of RIF-resistant TB were estimated in 2025 (7).

Resistance to RIF arises predominantly through mutations within specific regions of the *rpoB* gene, known as the RIF resistance-determining regions (RRDR). These include the N-terminal cluster, cluster I (amino acids 507–533), cluster II (amino acids 563–572) and cluster III (amino acid 687) according to *Escherichia coli* numbering (8, 9). Although first characterised in *Mycobacterium tuberculosis* (Mtb), *rpoB* mutations conferring rifampicin resistance have been reported in a broad and taxonomically diverse range of bacteria, including *Mycobacterium leprae*, *Staphylococcus aureus*, *Bacillus subtilis* and *Neisseria meningitidis* (10–13). Their broad taxonomic distribution, together with the clinical importance of rifampicin, has driven extensive research on *rpoB* mutations. This body of work has not only underpinned the development of rapid molecular diagnostics and surveillance for rifampicin-resistant pathogens but has also provided fundamental insights into bacterial structure and evolution. In structural biology, mapping RIF resistance mutations has helped define the architecture of the RNA polymerase active center, elucidate rifampicin binding, and reveal mechanisms governing transcription initiation, pausing, and termination (14). In evolutionary biology, RIF resistance has served as a model system for investigating the fitness costs of resistance, compensatory evolution, and adaptive trajectories that influence the persistence and spread of resistant lineages (15, 16).

Decades of research have progressively illuminated the molecular basis of antimicrobial resistance across diverse bacterial species, and this understanding has accelerated markedly with the advent of whole-genome sequencing (17–19). As a result, several specialized databases have emerged to catalog resistance-associated mutations, each built around a different scope, curation philosophy, and intended use case. The Comprehensive Antibiotic Resistance Database (CARD) is an ontology-centric knowledge base organized around the Antibiotic Resistance Ontology (ARO). It curates resistance gene sequences and resistance-conferring mutations across a broad range of pathogens. Recent expansions include likelihood-based mutations for *M. tuberculosis*, as well as data on resistance to disinfectants and antiseptics and on resistance-modifying adjuvants (20). The Drug Resistance Associated Genes database (DRAGdb) similarly provides curated resistance determinants, focusing on ESKAPE pathogens (i.e., *Enterococcus faecium*, *Staphylococcus aureus*, *Klebsiella pneumoniae*, *Acinetobacter baumannii*, *Pseudomonas aeruginosa* and *Enterobacter* spp.) together with *M. tuberculosis* (21). In contrast, the Comprehensive Resistance Prediction for Tuberculosis: an International Consortium (CRyPTIC) focuses on a single species, assembling the largest matched genotype–phenotype compendium available for Mtb (22). It includes whole-genome sequences and quantitative minimum inhibitory concentration (MIC) measurements for over 12000 clinical Mtb isolates across 13 antitubercular drugs, from which it derived a comprehensive catalog of resistance-conferring mutations (23). A third database, PointFinder, takes a narrower functional approach, targeting chromosomal point mutations specifically, with curated codon-level substitutions conferring resistance to quinolones, polymyxin, rifampicin, macrolides, and tetracyclines. It was initially developed around *Salmonella enterica*, *Escherichia coli*, and *Campylobacter jejuni* before extending to additional clinically relevant pathogens (24). Finally, the NCBI Reference Gene Catalog provides curated reference sequences for acquired resistance genes and species-specific point mutations across 31 drug classes and 58 drug phenotypes, alongside putative stress-response genes (acid, biocide, metal, and heat tolerance) and virulence genes—with particular depth for food-borne pathogens such as *E. coli* and *Salmonella* (25).

Despite the wealth of data, the current landscape of resistance resources for *rpoB* remains fragmented and difficult to navigate. Existing databases differ markedly in their taxonomic scope and curation standards. In addition, most retain each species’ native residue numbering, so the same *rpoB* position can refer to different amino acids across organisms. This convention precludes direct cross-species comparison and limits systematic integration among independent data sources. Moreover, most databases are designed to cover multiple antibiotics and resistance mechanisms simultaneously; RIF-specific data represent only a narrow subset of their overall content, resulting in sparse and incomplete coverage of *rpoB* rifampicin-associated resistance mutations.

To address these limitations, we developed RIFRMap (Rifampicin Resistance Mutation Map), an R Shiny web application for integrating, harmonizing, and interactively exploring RIF resistance-associated *rpoB* mutations. RIFRMap consolidates data from seven independent sources, covering clinical isolates, curated literature, and high-resolution experimental mutagenesis datasets, across 81 bacterial species, and standardizes mutation positions to a unified *E. coli* coordinate system to enable direct cross-species and cross-database comparisons. By presenting resistance-associated mutations through interactive visualizations rather than static tables, RIFRMap serves as a flexible platform for both education and research in antimicrobial resistance, allowing users ranging from researchers and clinicians to non-expert audiences to interrogate the biological patterns underlying RIF resistance.

## Materials and Methods

### Data sources and collection

RIFRMap integrates *rpoB* resistance mutation data from seven sources, whose taxonomic scope and update status are summarized in Table 1.

**Table 1.**
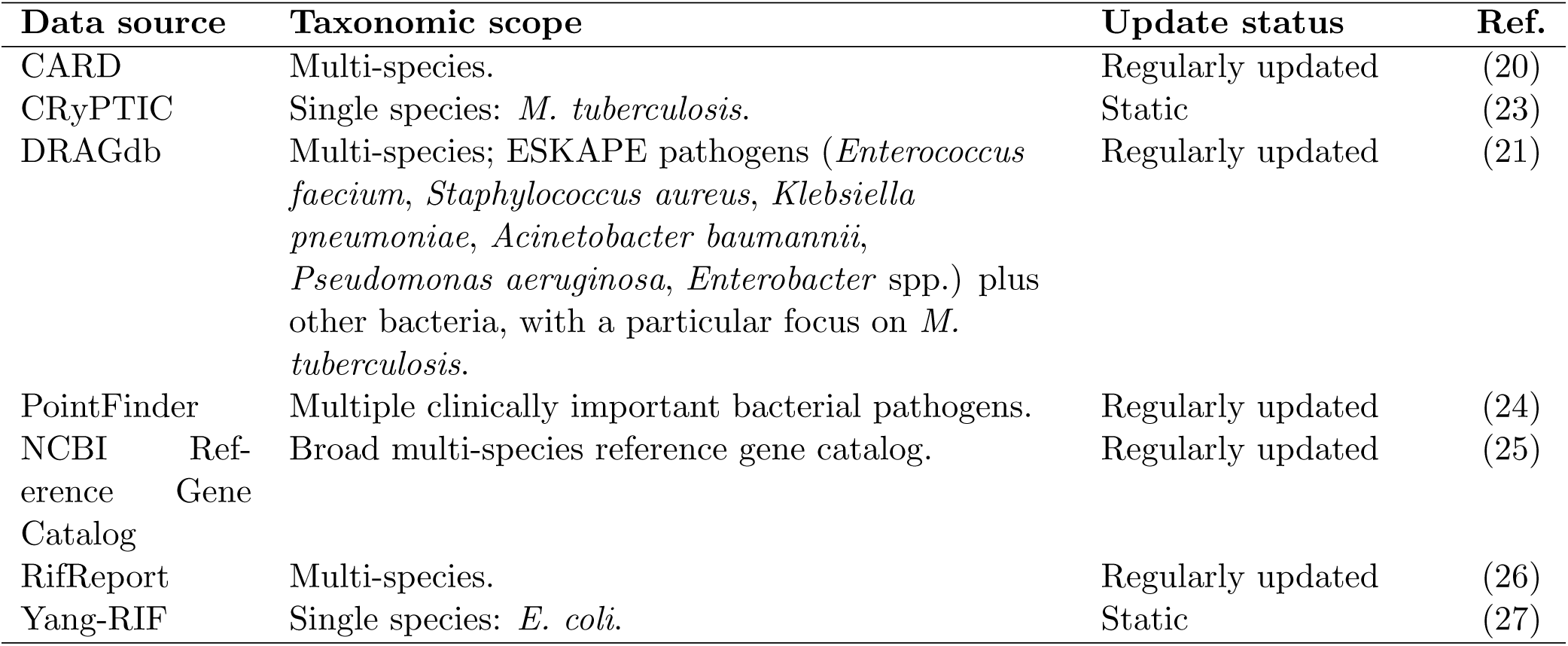
Mutation resources integrated into RIFRMap. Taxonomic scope, update status, and primary reference for each of the seven independent data sources consolidated by RIFRMap. Scope describes the range of bacterial taxa each resource covers; update status distinguishes resources under ongoing curation from those released as a fixed dataset.

| Data source | Taxonomic scope | Update status | Ref. |
|---|---|---|---|
| CARD | Multi-species. | Regularly updated | (20) |
| CRyPTIC | Single species: *M. tuberculosis*. | Static | (23) |
| DRAGdb | Multi-species; ESKAPE pathogens (*Enterococcus faecium*, *Staphylococcus aureus*, *Klebsiella pneumoniae*, *Acinetobacter baumannii*, *Pseudomonas aeruginosa*, *Enterobacter* spp.) plus other bacteria, with a particular focus on *M. tuberculosis*. | Regularly updated | (21) |
| PointFinder | Multiple clinically important bacterial pathogens. | Regularly updated | (24) |
| NCBI Ref-erence Gene Catalog | Broad multi-species reference gene catalog. | Regularly updated | (25) |
| RifReport | Multi-species. | Regularly updated | (26) |
| Yang-RIF | Single species: *E. coli*. | Static | (27) |

### CARD

The Comprehensive Antibiotic Resistance Database (CARD) is a manually curated repository of antibiotic resistance genes, mutations, and associated mechanisms (20). Each entry is drawn from peer-reviewed publications and reviewed by expert curators. For most species, CARD incorporates resistance variants based on direct experimental evidence. For *Mycobacterium tuberculosis*, where experimental antimicrobial susceptibility testing is particularly challenging, CARD employed a likelihood-based framework that infers resistance associations from single-nucleotide polymorphisms and observed phenotypic resistance across large collections of sequenced genomes. For RIFRMap, variant data were restricted to records of RIF resistance mediated by *rpoB*, classified under the single-resistance-variant parameter.

### CRyPTIC

The Comprehensive Resistance Prediction for Tuberculosis: an International Consortium (CRyPTIC) assembled a large-scale compendium of *M. tuberculosis* clinical isolates from 23 countries, pairing whole-genome sequencing with minimum inhibitory concentration (MIC) measurements for 13 anti-tuberculosis drugs (22). We downloaded all *rpoB* mutations for which RIF phenotypic data were available. Mutation-phenotype associations are derived from a genome-wide association study (GWAS), which employed interval regression in log_2_(MIC) to estimate the effect of each *rpoB* variant on RIF resistance (23). Positive values indicate increased resistance, whereas negative values denote increased susceptibility. False discovery rates were controlled using the Benjamini–Hochberg (BH) procedure. We retained only those entries constituting positive evidence of resistance, defined by a positive effect size (coefficient *>* 0) and a BH-corrected *p*-value *≤* 0.05.

### DRAGdb

The Drug Resistance Associated Genes database (DRAGdb) is a manually curated database of antibiotic resistance mutations that includes ESKAPE pathogens (i.e., *Enterococcus faecium*, *Staphylococcus aureus*, *Klebsiella pneumoniae*, *Acinetobacter baumannii*, *Pseudomonas aeruginosa* and *Enterobacter* spp.) as well as other bacteria, with a special focus on *M. tuberculosis* (21). All *rpoB* mutations associated with RIF resistance were retrieved along with their corresponding Protein Variation Effect Analyzer (PROVEAN) scores (28). PROVEAN predicts the functional impact of amino acid substitutions based on evolutionary sequence conservation. Mutations with scores *≤ −*2.5 are classified as deleterious—suggesting a disruption of protein function—while scores above this threshold are considered neutral. To isolate substitutions with a probable functional impact on the RNA polymerase *β* subunit, we restricted our dataset exclusively to these deleterious mutations.

### PointFinder

PointFinder, developed at the Center for Genomic Epidemiology, is a web tool designed to identify chromosomal point mutations associated with antimicrobial resistance (24). It curates point mutations in key resistance genes in a defined set of clinically significant bacterial species, with entries supported by peer-reviewed literature. The PointFinder data were filtered to retain only *rpoB* point mutations associated with RIF resistance.

### NCBI Reference Gene Catalog

The NCBI Reference Gene Catalog is a curated reference database of bacterial genes and point mutations associated with antimicrobial resistance. It forms the core component of the AMRFinderPlus database, along with the Reference Gene Hierarchy and the Reference HMM Catalog, and provides the reference data underlying the AMRFinderPlus software for genomic antimicrobial resistance screening (25). We extracted all entries in which the annotation of the gene family was *rpoB* and the drug subclass included Rifampin.

### RifReport

The curated dataset was originally assembled by Bolourchi *et al.* through a systematic literature survey to compile a comprehensive catalog of reported rifampicin resistance-associated *rpoB* mutations (26). Studies were identified through Google Scholar using the following keywords: rifampicin resistance, RNA polymerase subunit beta, point mutation, and amino acid substitution. The dataset is maintained as a continuously updated repository in https://github.com/JanEngelstaedter/ResistanceSurveys, from which we retrieve the most current version.

### Yang-RIF

Yang *et al.* used multiplex automated genome engineering to generate a saturated pool of all 760 variants at the complete RIF binding site of *Escherichia coli* RNA polymerase (positions 510–537 and 563–572) (27). Drug-specific effects were quantified by normalizing mutant abundances after RIF treatment to mutant abundances under untreated growth conditions, yielding logarithmic fold-change values for each substitution. Substitutions with log_2_ FC *≥* 0 were retained as being enriched under RIF selection.

### Data processing

Next, for each bacterial species represented in the integrated dataset, a single reference strain was assigned, and its genome was retrieved from the NCBI Genome database (29). The *rpoB* sequence of each genome was extracted from their annotated genomic coding sequences using the following keywords: rpoB, DNA-directed RNA polymerase subunit beta, DNA-directed RNA polymerase subunit beta chain, and RNA polymerase, beta subunit. To standardize mutation positions for cross-species comparison, the sequences were translated and aligned with the *rpoB* reference sequence of *E. coli* MG1655. The resulting alignment was then utilized to map each species-specific position to its corresponding *E. coli* coordinate.

### Implementation

RIFRMap was implemented in R (version 4.5.3) (https://www.R-project.org) using Shiny (version 1.13.0) (https://CRAN.R-project.org/package=shiny) and hosted on https://rifrmap.org. Bioinformatics analyses were performed using the packages rentrez v1.2.4 (https://CRAN.R-project.org/package=rentrez), ALJEbinf v0.1.0 (https://github.com/JanEngelstaedter/ALJEbinf), Biostrings v2.78.0 (https://bioconductor.org/packages/Biostrings) and grantham v0.1.4 (https://CRAN.R-project.org/package=grantham). Data processing and manipulation were conducted with tidyverse v2.0.0 (https://CRAN.R-project.org/package=tidyverse). Visualization and interactivity are implemented using ggplot2 v4.0.3 (https://CRAN.R-project.org/package=ggplot2), ggiraph v0.9.6 (https://CRAN.R-project.org/package=ggiraph), and ggupset v0.4.1 (https://CRAN.R-project.org/package=ggupset). Summary tables were produced using gtsummary v2.6.1 (https://CRAN.R-project.org/package=gtsummary) and rendered with gt v1.4.0 (https://CRAN.R-project.org/package=gt). The source code is publicly available at https://github.com/lna0104/RIFRMap.

## Results and discussion

RIFRMap is an interactive web application for integrating, harmonizing, and visualizing RIF resistance-associated *rpoB* mutations across bacteria. The design of RIFRMap includes five pages: *Home*, *Explore*, *Data*, *Docs*, and *Team*. The *Home* page provides an overview of the application and its data sources (Figure 1A), while the *Docs* page presents detailed descriptions of each data source, including brief explanations of their content and origin (Figure 1B). The *Data* page provides full access to the integrated dataset through two searchable tables: the mutation data table catalogs all resistance-associated mutation entries, while the reference table provides complete bibliographic details, including DOI and PubMed ID, allowing users to trace each mutation to its original publication (Figure 1C). Both tables support full-text search, column sorting, and csv export. The *Team* page lists project contributors, institutional affiliations, and ORCIDs, alongside contact details for user feedback.

**Figure 1.**
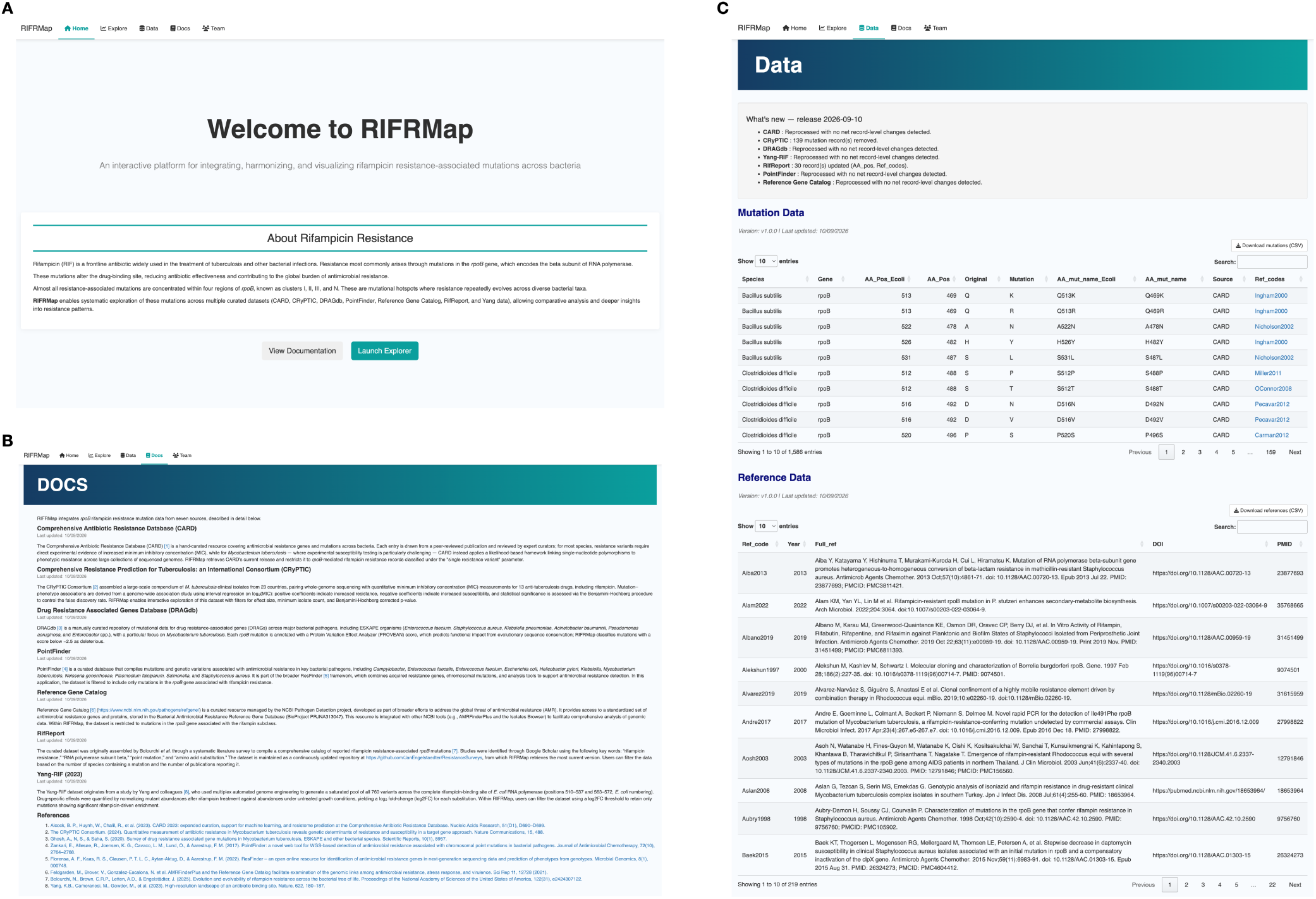
Overview of the RIFRMap user interface. (A) The *Home* landing page. The *Home* page provides an overview of our application, featuring integrated navigation gateways that direct users to the interactive *Explore* and *Docs* pages. (B) The *Docs* page documents each of the seven integrated sources, detailing scope, curation methodology, and RIFRMap inclusion criteria alongside primary literature citations. (C) The *Data* page. The mutation data table contains 1586 entries and displays each resistance-associated mutation with species, gene, *E. coli* -mapped amino acid position, species-specific amino acid position, reference amino acid, mutant amino acid, standardized mutation name in *E. coli* coordinates, species-specific mutation name, data source, and linked reference codes. The reference table contains 219 entries and provides bibliographic information for the primary references cited across the seven integrated sources, including reference code, year, full citation, DOI, and PubMed ID (PMID). Both tables support csv export for downstream use.

The *Explore* page comprises six modules: *Data Filters*, *Mutation Map*, *Mutation Frequency*, *Dataset Comparison*, *Summary* and *Filtered Data*. Together, these modules enable users to apply granular data filtering and interactively investigate mutation patterns, frequency distributions, and cross-dataset relationships. All graphics outputs can be downloaded in multiple formats (png, pdf, jpg, and tiff) for storage, publication, or further analysis.

### Overview of Integrated Data

At the time of publication, RIFRMap aggregates a total of 1586 entries from seven sources, representing 394 unique mutations at 114 *E. coli* -mapped positions, reported across 81 bacterial species. The RifReport dataset contributes the largest number of mutations (220) and the broadest taxonomic coverage, spanning 72 species (Figure 2). While most sources include data for multiple taxa, the CRyPTIC and Yang-RIF datasets are restricted to single species *M. tuberculosis* and *E. coli*, respectively (Figure 2B). Nevertheless, these sources provide the highest mutational data density for their respective target organisms. *M. tuberculosis* is the most widely represented species, with 183 variants compiled across CRyPTIC, CARD, DRAGdb, PointFinder, and RifReport (Figure 2C). *E. coli* is the second most frequent organism (160 mutations), driven primarily by the Yang-RIF experimental dataset together with RifReport. Beyond these two organisms, representation drops markedly, with most species contributed by RifReport alone, underscoring the relatively limited systematic characterization of *rpoB* mutations in non-TB pathogens.

**Figure 2.**
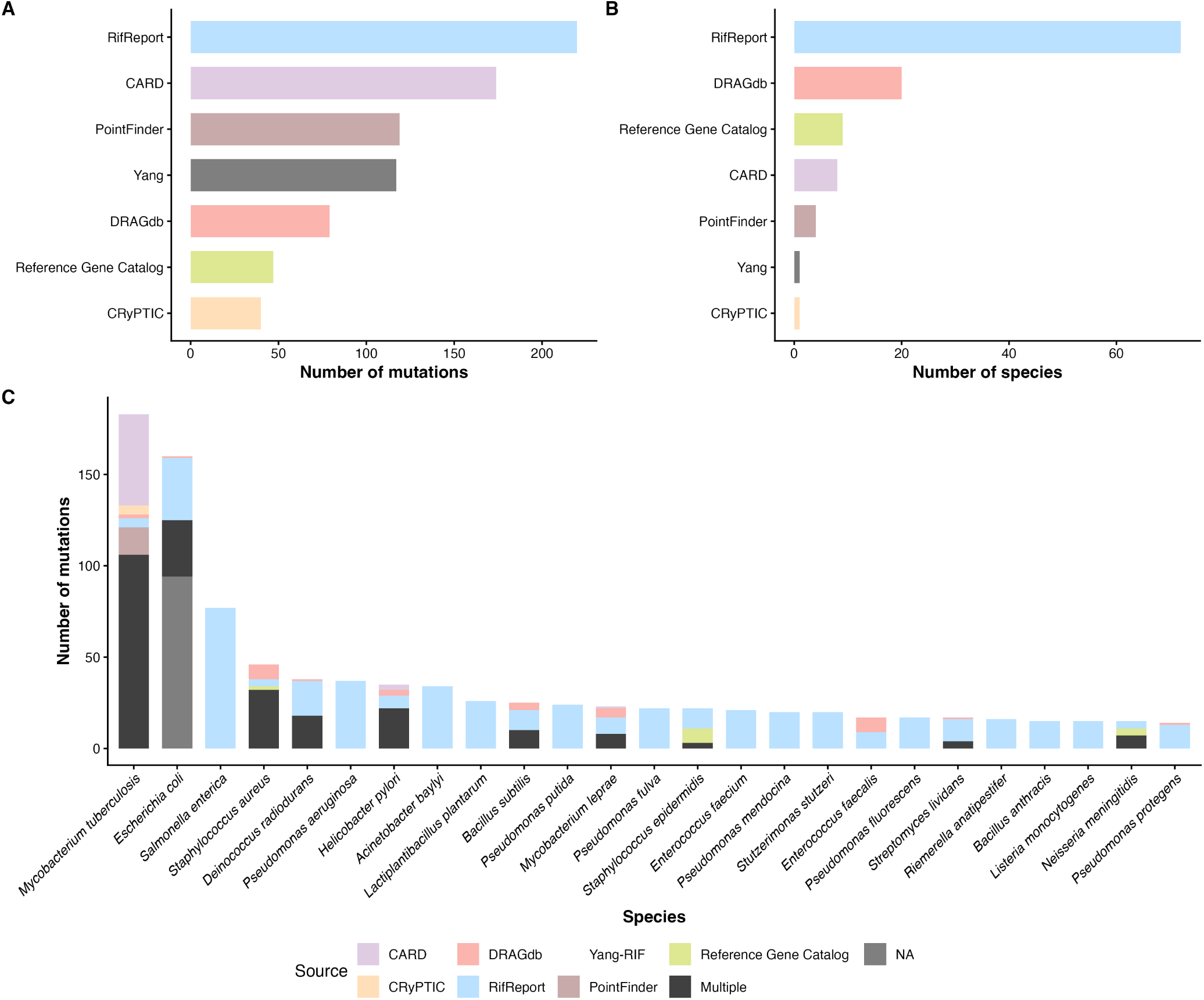
Overview of integrated data sources. (A) Number of mutations contributed by each of the seven integrated data sources, where a mutation is identified by its *E. coli* -mapped position and mutant residue. (B) Number of bacterial species represented within each data source. (C) Total number of unique mutations reported for the top 25 bacterial species across all integrated datasets, stacked by contributing source. Segments drawn in dark gray and labeled “Multiple” denote mutations reported for a given species by more than one source. Panels A and B are color-coded by data source, whereas panel C additionally includes the ”Multiple” category.

### Module 1 - Data Filters

The *Data Filters* module within the *Explore* page serves as the primary control interface through which users can define the scope of all downstream visualizations (Figure 3). To initiate a session, users first select which of the seven data sources to include. For multi-species datasets, an additional species-level control further restricts the selected data to one or more organisms of interest. Beyond these universal filters, source-specific parameters allow for granular refinement of the mutation dataset. For CRyPTIC-derived data, users can apply thresholds on GWAS effect size (log_2_(MIC) coefficient), the minimum and maximum number of isolates carrying a mutation, and Benjamini–Hochberg-corrected *p*-value. DRAGdb entries can be filtered by PROVEAN scores to prioritize variants predicted to have strong functional impact. RifReport entries can be subset by the number of supporting studies and the number of species in which a mutation has been reported, enabling users to prioritize mutations supported by broader evidence. Yang-RIF mutations can be refined using a log_2_(fold-change) threshold to retain only mutations that exhibit positive RIF-selected enrichment. All filters operate simultaneously, updating all downstream visualizations upon application, and can be reset to their defaults at any time.

**Figure 3.**
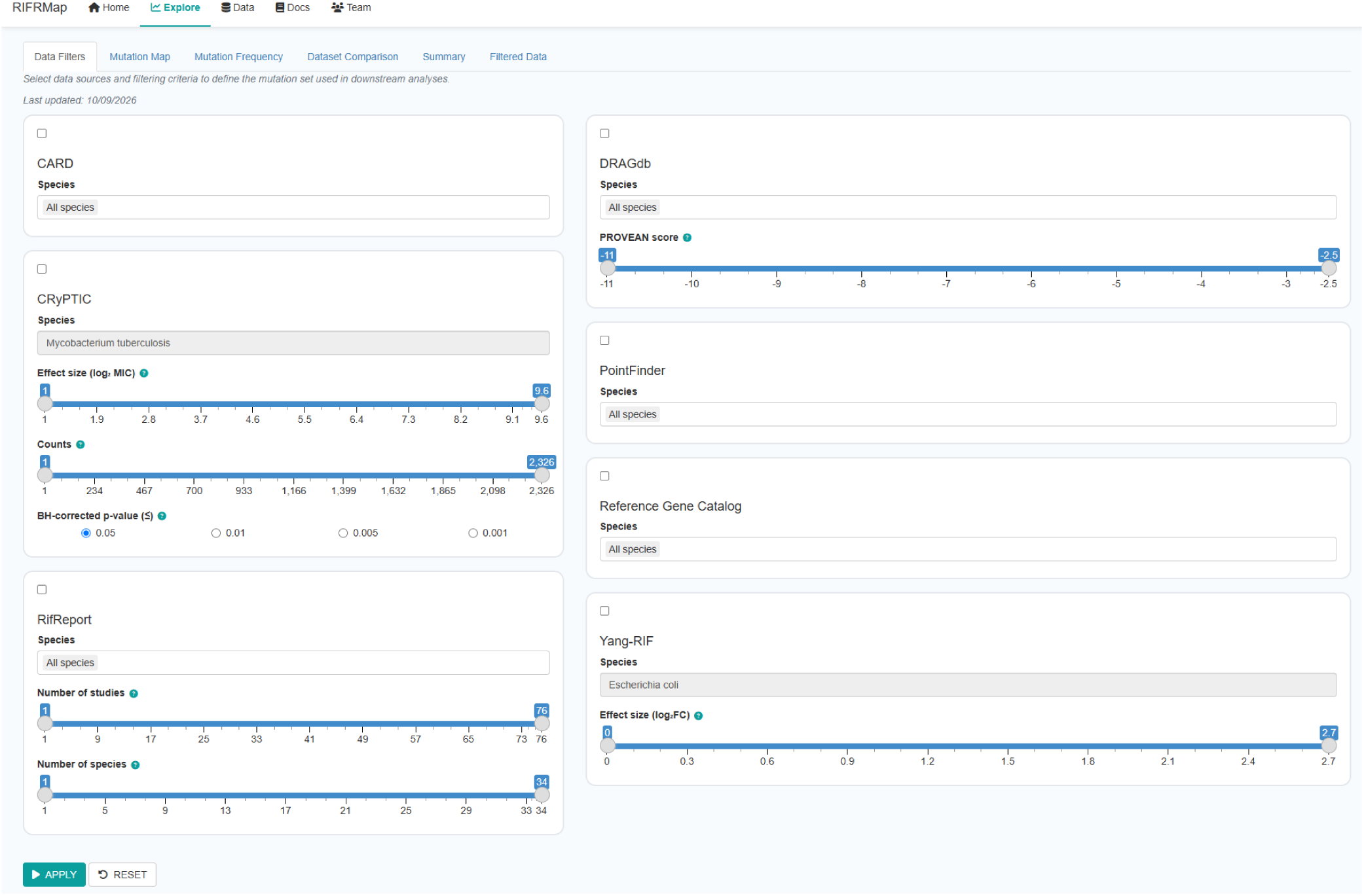
The *Data Filters* module on the *Explore* page, which serves as the primary control interface for defining the scope of all downstream visualizations.

### Module 2 - Mutation Map

The *Mutation Map* module provides a comprehensive, species-centric grid overview of the distribution of RIF resistance-associated mutations in the *rpoB* gene. By default, the plot displays species-specific positions, reference amino acids, and observed mutant residues across the major clusters of the RIF binding site (clusters N, I, II, and III). Users can also select an all-mutations view, which displays all mutations, including those falling outside the four defined clusters. In both views, position numbers are shown in species-specific coordinates, reflecting the native numbering system of each organism, while the underlying layout is organized according to *E. coli* -mapped positions, enabling direct visual cross-species comparison. In addition, mutant amino acids can be color-coded according to three user-selectable schemes: distribution across species, physicochemical type of amino acid, or Grantham distance between reference and mutant residues (30).

Figure 4 illustrates a representative use case of the *Mutation Map* module when all data sources are activated. Across all species, resistance mutations are predominantly concentrated in cluster I (positions 507–533). Within this region, position 526 (*E. coli* numbering) emerges as the most mutationally diverse site, harboring substitutions to all nineteen alternative amino acids. The breadth of substitutions at this position is largely attributable to the Yang *et al.* saturation mutagenesis dataset, which systematically generated and screened all possible amino acid substitutions across the RIF binding site (27).

**Figure 4.**
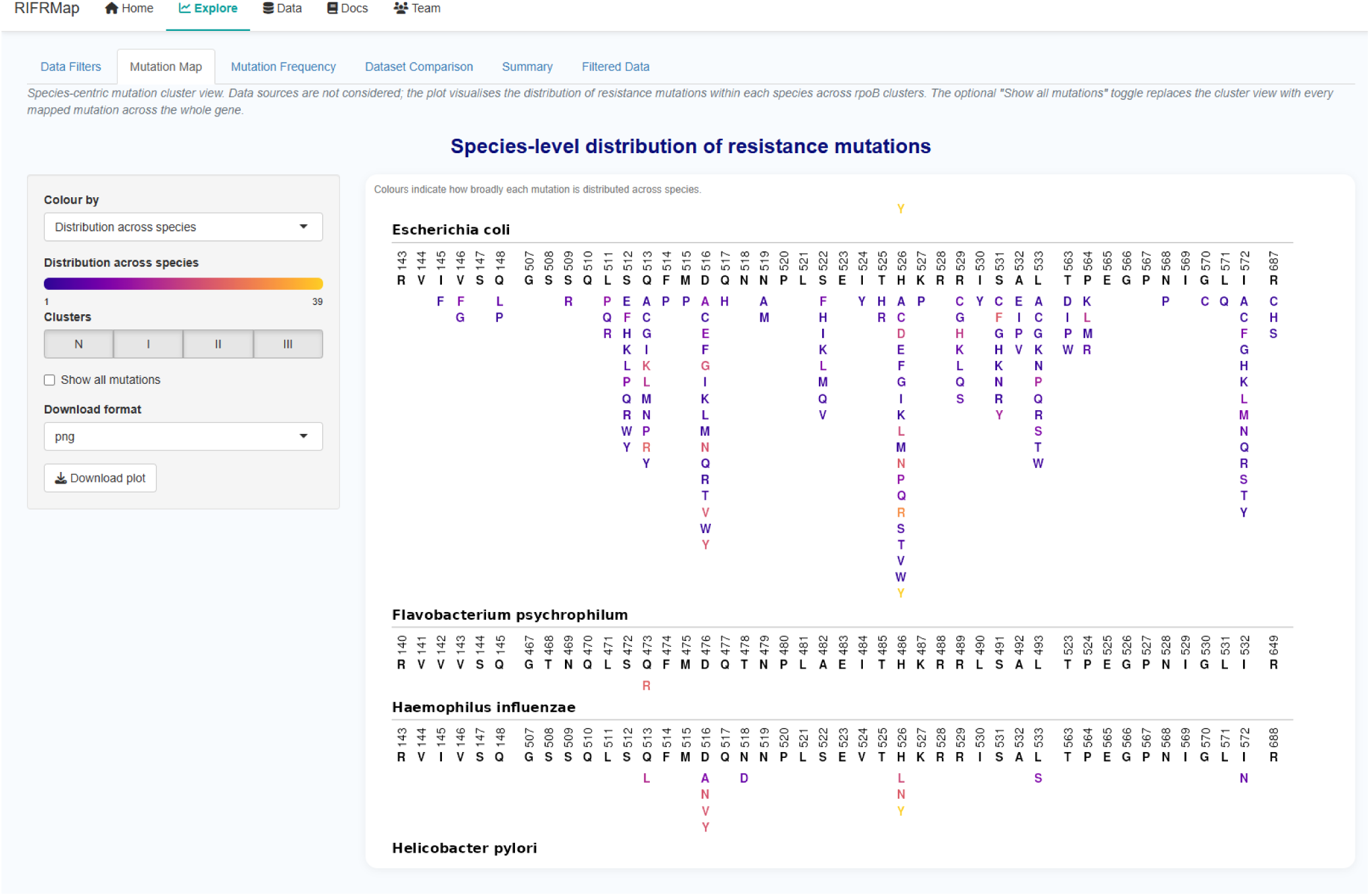
The *Mutation Map* module, showing the species-level distribution of RIF resistance mutations across *rpoB* with all data sources activated. Each block corresponds to a single bacterial species, with rows displaying the position number, the reference amino acid (shown in black), and all observed mutant amino acids. Navigational controls allow the user to toggle between the major functional regions of the RIF binding site (clusters N, I, II, and III) or view all reported mutations across the gene. Mutant amino acids are color-coded based on their distribution across the represented species.

### Module 3 - Mutation Frequency

Whereas the *Mutation Map* provides a species-centric view, the *Mutation Frequency* module offers a position-centric perspective, revealing how broadly each mutation is distributed across all species in the filtered dataset. The module features an interactive heatmap whose rows are the twenty standard amino acids and whose columns are the *E. coli* -mapped *rpoB* positions at which at least one mutation is recorded (Figure 5). Each cell is color-coded by the number of species in the filtered dataset that harbor the corresponding substitution, while white cells denote unobserved variants. Selecting any cell populates a companion table detailing all species in which that substitution has been reported. Figure 5 presents the complete *rpoB* RIF resistance landscape across all integrated sources. Mutations are predominantly concentrated within cluster I; although positions 507–533 span just 27 of the 1342 residues (2.0% of the gene), they account for 1231 of the 1586 records (78%) and 232 of the 394 unique mutations (59%) in the integrated dataset. Notably, the mutation 526Y is the most frequently observed variant, occurring across 39 species. In contrast, outside cluster I—particularly in regions spanning positions 25–506, 537–562, and 583–1314—most mutations are infrequent and typically restricted to single species. This could be due to lineage-specific variation rather than broadly conserved resistance mechanisms, or it could reflect ascertainment biases (given that many studies focus on cluster I).

**Figure 5.**
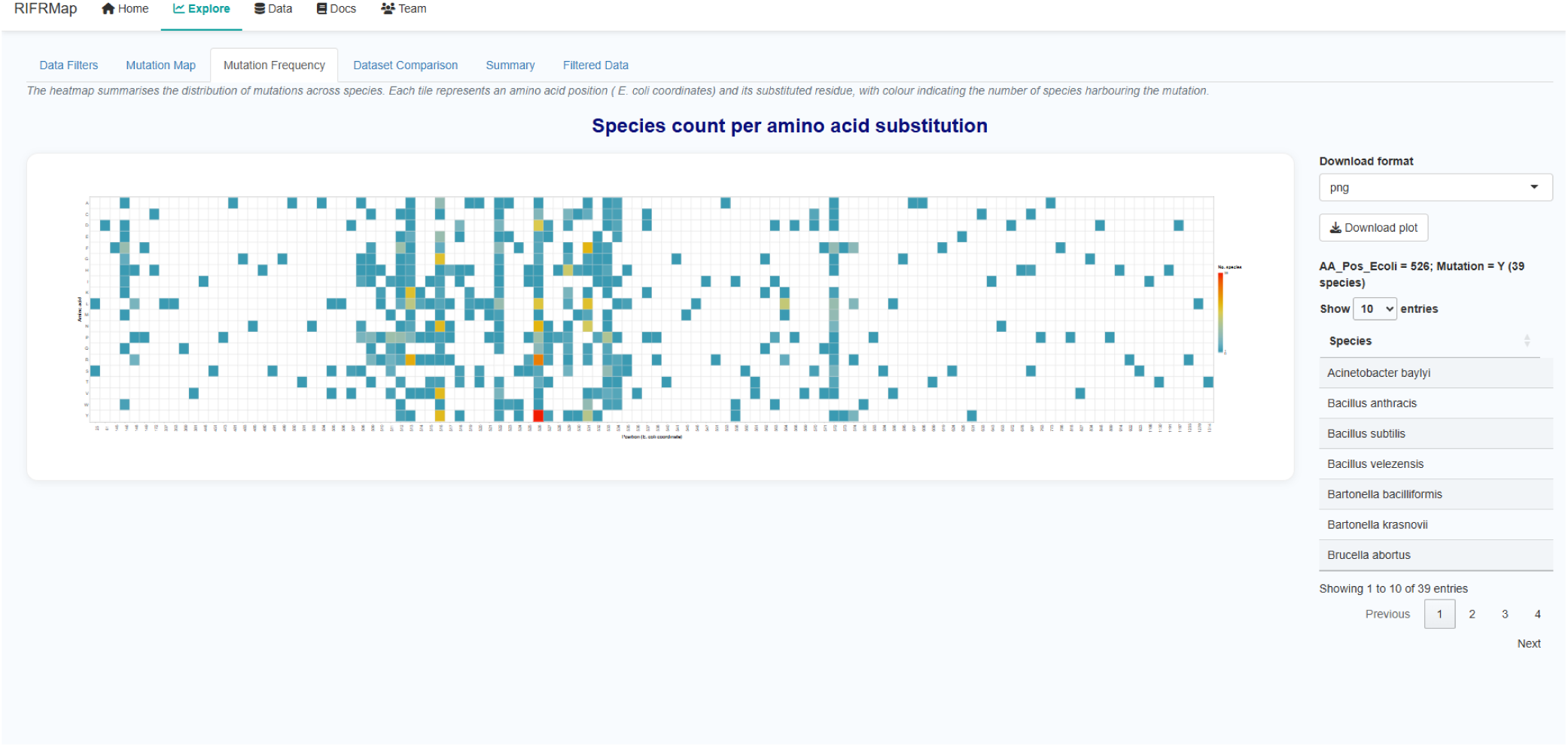
The *Mutation Frequency* module, shown with all data sources activated. The interactive heatmap carries one row for each of the twenty standard amino acids and one column for each *E. coli* -mapped *rpoB* position at which a mutation is recorded. Each tile represents a single mutation. Tile color denotes the number of species in which that substitution has been reported, ranging from blue (low prevalence) to red (high prevalence); white cells denote unobserved variants. The substitution 526Y (highlighted in red) is the most broadly distributed RIF resistance variant in the integrated dataset, reported in 39 species. The companion table on the right populates when you select a cell, showing all species carrying the selected substitution.

### Module 4 - Dataset Comparison

When integrating data from multiple resistance databases, assessing the degree of agreement among sources becomes essential: convergent evidence across distinct sources provides stronger empirical validation for a given mutation, while source-exclusive variants may warrant more cautious interpretation. The *Dataset Comparison* module addresses this need through two complementary visualizations.

Figure 6 presents a representative example of this module with all integrated data sources enabled. Panel A presents an upset-style bar graph quantifying the number of mutations exclusive to each source and the sizes of their intersections. RifReport contributed the most source-exclusive mutations (86 of 220), reflecting its broad taxonomic scope and systematic assembly from published literature. Yang-RIF contributed the second largest number of unique mutations (52 of 117), reflecting the comprehensive experimental coverage of the saturation mutagenesis approach, many of which have never been observed in clinical or environmental isolates. By contrast, the CRyPTIC dataset and the NCBI Reference Gene Catalog contribute the fewest source-exclusive mutations, indicating substantial overlap with other databases.

**Figure 6.**
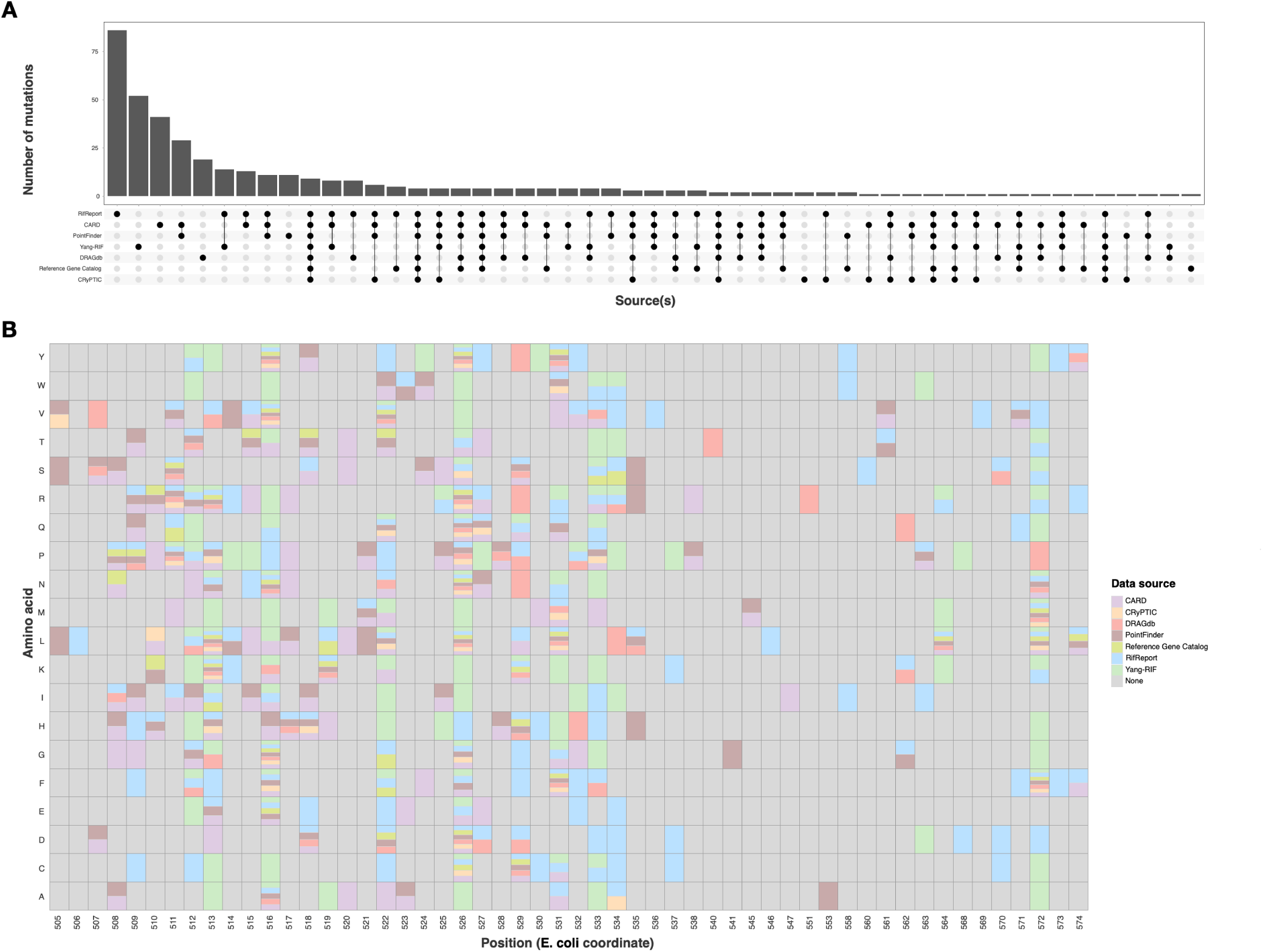
The *Dataset Comparison* module, shown with all data sources activated. (A) An upset-style intersection plot in which each bar counts the mutations reported by the combination of sources marked beneath it. Bars marked with a single dot therefore give the mutations exclusive to that source, and bars spanning several dots show the sizes of the corresponding intersections. RifReport provides the largest proportion of source-exclusive variants, reflecting its extensive taxonomic and historical scope. (B) A high-resolution source-attribution heatmap mapping *rpoB* positions (*E. coli* numbering) against the twenty standard amino acids. Each tile is partitioned into proportional, color-coded segments, where each color corresponds to a specific contributing data source. Gray shading denotes individual position-residue combinations for which no resistance-associated mutation was reported. For clarity, only the high-density region spanning positions 505–574 is displayed.

Panel B presents a source-attribution heatmap in which rows correspond to the twenty standard amino acids and columns to *E. coli* -mapped *rpoB* positions. When a variant is reported by multiple independent sources, the cell is divided into proportional color-coded segments representing each contributing database. Moreover, the *Dataset Comparison* module allows users to show or hide individual sources through the legend controls. This visualization helps users to immediately distinguish mutations exclusive to a single study from those robustly corroborated across independent datasets.

Cross-source corroboration is concentrated in the same regions that dominate the mutation landscape as a whole: of the 182 mutations reported by two or more sources, 149 (82%) fall within cluster I (positions 507–533) or cluster II (positions 563–572) (Figure 6B). Nine mutations are reported simultaneously across all seven integrated sources: 513K, 513L, 516G, 516V, 516Y, 526L, 526N, 526Y, and 572F. These variants represent the most extensively validated resistance markers. Outside positions 505–574, by contrast, 95% of mutations are reported by only one or two sources.

### Module 5 – Summary

The *Summary* module provides a quantitative overview of the user-selected dataset, complementing the three preceding visual modules with tabulated counts and distributional summaries. It comprises three elements, all recomputed whenever a new filter selection is applied. A summary table reports the composition of the filtered dataset, giving the number and proportion of records contributed by each bacterial species, gene, and data source. Alongside it, a bar chart ranks the fifteen most frequently reported amino-acid substitutions (named by their *E. coli* -mapped position), and a histogram shows how records are distributed across *E. coli* -mapped *rpoB* positions. All three elements are computed over records rather than over unique mutations: a substitution reported by multiple sources, or observed across multiple species, contributes one count per report. By quantifying reporting frequency rather than taxonomic breadth—the latter being the focus of the *Mutation Frequency* module—the *Summary* module offers a complementary perspective on the overall weight of evidence supporting a given variant.

### Module 6 – Filtered Data

The *Filtered Data* module gives direct access to the individual records underlying the *Summary* module’s statistics. It lists every record retained by the current filter settings, giving each record’s species, gene, native and *E. coli* -mapped position and mutation codes, data source, and linked reference code(s); as in the *Summary* module, a record is counted once per report, so a substitution observed across multiple sources or species appears as multiple rows. The table can be exported as a csv file for further analysis, complementing the unfiltered Data tab by giving users the subset matching their chosen filters rather than the full dataset.

## Conclusions

RIFRMap integrates resistance-associated *rpoB* entries from seven independent sources into a unified, interactively explorable platform. By standardizing heterogeneous datasets to an *E. coli* coordinate system and presenting them through a suite of complementary visualizations, RIFRMap substantially lowers the barrier to cross-species and cross-dataset comparison. Analysis across modules confirms that resistance mutations are predominantly concentrated within cluster I of the RRDR, with position 526 emerging as the most mutationally diverse site and the 526Y substitution exhibiting the greatest taxonomic breadth. Furthermore, by integrating saturation mutagenesis data alongside clinical and curated repositories, RIFRMap reveals the gap between theoretically accessible and naturally observed resistance variants—a distinction with implications for surveillance and experimental design. While the current version is limited to point mutations in *rpoB*, future development may incorporate indel variants, additional resistance genes, phenotypic effects of resistance mutations, and phylogenetic context. We anticipate that RIFRMap will serve as a long-term community resource for researchers, clinicians, and educators working at the intersection of antimicrobial resistance, bacterial genomics, and evolutionary biology.

## Acknowledgements

We are grateful to Negin Bolourchi for the systematic literature survey that produced the RifReport dataset and for the continued work curating and expanding it.

## Notes

### Competing Interest Statement

The authors have declared no competing interest.

